# Error-Related Memory Biases Are Specific to Social Stimuli for Socially Anxious Individuals

**DOI:** 10.1101/2025.10.13.682081

**Authors:** Kianoosh Hosseini, Aaron T. Mattfeld, Jeremy W. Pettit, George A. Buzzell

## Abstract

Social anxiety (SA) is associated with enhanced error monitoring, yet underlying mechanisms remain unclear. Consistent with cognitive models of SA, we propose that stronger error monitoring contributes to SA by strengthening memory encoding of errors (including relevant social cues), negatively biasing what is remembered. Supporting this hypothesis, our prior work demonstrated that high SA individuals exhibit better memory for faces presented during error (vs. correct) trials. To test whether this Memory Bias for Error Events is specific to social stimuli, 140 participants completed a Flanker task with trial-unique faces (social) or objects (non-social) as background images, followed by a surprise memory test. Results revealed that higher SA symptoms were associated with enhanced memory for faces on error (vs. correct) trials, but not for objects. These findings replicate and extend our prior work, demonstrating that SA-related memory biases for errors are specific to social stimuli, rather than reflecting general encoding biases.

## Introduction

Social anxiety (SA) is a highly prevalent and debilitating condition characterized by an intense fear of social interactions that significantly impairs interpersonal relationships and occupational functioning (American Psychiatric Association, 2022; Kessler et al., 2005; Morrison & Heimberg, 2013). Despite the high prevalence of SA, treatment effects remain modest (Carpenter et al., 2018; Hoyer et al., 2016), underscoring the need to identify mechanistic targets. Research shows SA and related symptoms are associated with enhanced error monitoring (Barker et al., 2015; Endrass et al., 2014; Hosseini et al., 2024; Judah et al., 2016; Kujawa et al., 2016; Meyer et al., 2021; Niu et al., 2023), which refers to the detection and processing of self-committed errors (Gehring et al., 1993). However, the causal nature of associations between SA symptoms and error monitoring remains unclear. In line with cognitive models of SA (Clark & Wells, 1995; Rapee & Heimberg, 1997), we have proposed that the link between error monitoring and SA may be cyclical: enhanced error monitoring in SA drives memory biases for error events, strengthening negative beliefs about one’s abilities and ultimately maintaining or worsening SA (Hosseini et al., 2024). Our prior study provided initial evidence by linking greater SA symptoms to an increased memory bias for error events (Hosseini et al., 2024). However, our previous work was limited in terms of sample size and did not test whether the memory bias was specific to social stimuli for high SA individuals. In an effort to advance theory and inform the design of novel treatments for SA, the current study sought to close these gaps.

Cognitive models of SA propose that individuals high in SA symptoms excessively monitor their behavior and internal states during social interactions, motivated by an underlying fear of negative evaluations (Clark & Wells, 1995; Rapee & Heimberg, 1997). According to these models, heightened self-focused attention and self-monitoring impacts what is processed and encoded into memory during social events, leading to memory biases that contribute to negative self-appraisals that ultimately maintain or worsen SA symptoms. Supporting these ideas, studies employing neural measures have shown that individuals high in SA exhibit enhanced error monitoring—an aspect of self-monitoring localized to the medial frontal cortex (Barker et al., 2015; Buzzell et al., 2017; Endrass et al., 2014; Hosseini et al., 2024; Judah et al., 2016; Kujawa et al., 2016; Meyer et al., 2021; Niu et al., 2023). Increased error monitoring likely directs attention towards error events (Maier et al., 2011; Steinhauser & Andersen, 2019; Wessel, 2018), increasing the salience of error-related information (Hajcak et al., 2005). As we have previously argued (Hosseini et al., 2024) heightened attentional orienting towards errors could lead to enhanced encoding of error events, producing a subsequent memory bias that favors error-related information over information associated with correct responses. This proposed mechanism is consistent with research demonstrating that individuals high in SA exhibit memory biases for negative aspects of their performance (Cody & Teachman, 2010; Romano et al., 2020).

Linking prior work on enhanced error monitoring and memory biases in SA, we recently provided the first empirical evidence demonstrating that individuals high in SA symptoms exhibit an enhanced Memory Bias for Error Events (Hosseini et al., 2024). In our prior study, we had participants complete a social-evaluative cognitive control task (a Flanker task under peer observation) with trial-unique social stimuli (faces) presented on every trial. Following the task, participants then completed a surprise memory assessment to probe their incidental memory for the faces that originally appeared on error vs. correct trials. Using this paradigm, we found that individuals high in SA exhibited better memory for faces that appeared on error (vs. correct) trials, which we refer to as Memory Bias for Error Events. Furthermore, we found neural activity associated with error monitoring was not only enhanced for individuals with greater SA symptoms, but it also directly predicted Memory Bias for Error Events. However, as our prior study only employed faces as background images, it remains unknown whether individuals high in SA preferentially encode any information present during error events, or whether their error-related encoding is specifically biased toward socially-relevant information.

We hypothesize that an enhanced Memory Bias for Error Events in high SA individuals is specific to social stimuli. As described below, this hypothesis is based on the understanding that salient events (like errors) trigger biased processing and encoding of motivationally relevant information, and that for individuals high in SA, social stimuli are motivationally relevant. Errors reflect a specific instance of a broader class of “salient events” (i.e., unexpected and/or high-valenced events) which trigger norepinephrine (NE) release and a rapid increase in attention and arousal (Dahl et al., 2022; De Martino et al., 2008; Mather et al., 2016; Ullsperger et al., 2010; Wessel, 2018; although evidence of NE release for errors is indirect). Literature on how other kinds of salient events impact memory suggests that this heightened arousal state facilitates preferential encoding of stimuli/information concurrently present and motivationally relevant to the individual (Clewett et al., 2018; Dunsmoor et al., 2015; Hutmacher & Kuhbandner, 2020; Lin et al., 2010; Patil et al., 2017; Swallow et al., 2022; Swallow & Jiang, 2010, 2011). Briefly, this is thought to occur due to NE release driving an increase in neural activity within brain areas that are concurrently active (due to their processing of information that is either task relevant or otherwise motivationally relevant) and suppression of other neural activity (see Mather et al., 2016 and Mather & Sutherland, 2011 for further details). In other words, salient events trigger biased attention towards motivationally-relevant stimuli, and as a result, yield biased encoding of those stimuli. For individuals high in SA, whose primary fear is negative evaluation *by others* (Clark & Wells, 1995; Rapee & Heimberg, 1997; Zhang et al., 2023), social cues like faces are highly motivationally relevant, demonstrated by biased processing of such stimuli in SA (Cooney et al., 2006; Kivity & Huppert, 2016). Moreover, we anticipate the motivational relevance of social stimuli would be particularly prominent during error commission, as this reflects an instance of performance failure. Thus, we hypothesize that when individuals high in SA make errors, they will exhibit enhanced processing of social stimuli (e.g., faces) as opposed to other, non-social stimuli that might be present during an error (e.g., inanimate objects), leading to a Memory Bias for Error Events that is selective for social stimuli. This hypothesis is also consistent with other work demonstrating that other kinds of negative memory biases in SA are specific to social-evaluative information (Romano et al., 2020).

The present study was designed to address two primary aims, which build upon our prior findings. Our first aim was to replicate the finding that higher SA symptoms are associated with a heightened Memory Bias for Error Events, doing so within the context of a larger, independent sample and while employing a more robust memory assessment approach. Specifically, we replaced the “old/new” recognition task used in our prior study with a two-alternative forced choice (2AFC) recognition paradigm. Use of a 2AFC paradigm requires participants to make relative (as opposed to absolute) memory judgments and is therefore less susceptible to response biases (i.e., individual differences in the overall tendency to respond “old” vs. “new”) that could potentially confound memory effects (Kellen et al., 2018; Kroll et al., 2002; Van Zandt, 2000). Our second aim was to determine whether a heightened Memory Bias for Error Events in high SA individuals is specific to socially-relevant stimuli present during error events, or if this effect generalizes to non-social stimuli. To address both aims, we employed a between-subjects design in which 140 participants completed a modified Flanker task featuring either trial-unique face (social stimuli) or inanimate object (non-social stimuli) as background images, followed by a surprise 2AFC recognition memory task. If the memory bias observed in individuals high in SA reflects a general tendency to encode all information present at the time of error events, then an enhanced Memory Bias for Error Events should be observed regardless of whether background images are social or non-social in nature. However, if the memory bias is specific to socially-relevant information, then an enhanced Memory Bias for Error Events among high SA individuals should only emerge among participants that complete the Flanker task employing face images. Understanding whether such memory biases are socially specific would advance SA theory and could inform novel treatment designs that target selective encoding of social information during error events rather than addressing error monitoring or memory biases broadly.

## Transparency and Openness

### Preregistration

This experiment was not preregistered.

### Data, materials, code, and online resources

The PsychoPy tasks, questionnaires, the inanimate image dataset, and data analysis scripts are publicly available on the following GitHub repository: https://github.com/NDCLab/mfe-c-dataset. Deidentified data are available from the corresponding author upon request.

### Reporting

We report how we determined our sample size, all data exclusions, all manipulations, and all relevant measures in the study.

### Ethics approval

Ethics approval was granted by the Florida International University (FIU) Social and Behavioral Institutional Review Board (Protocol number: 110925) for this study.

## Methods

### Participants

An a priori power analysis conducted in G*Power (Faul et al., 2009) determined that 100 participants were needed to achieve 80% power (α = 0.05) to detect a small-to-medium effect size (f² = 0.1) for the primary analysis of interest (R² increase reflecting a significant interaction for a multiple linear regression model). Therefore, to allow for potential data exclusion, we set our a priori sample size (stopping rule) at N=140. To this end, participants included in this study were 140 healthy adults (M = 22.42 years, SD = 3.31; 113 female, 27 male; see Supplemental Table S6 for detailed participant demographics) with no history of head injuries causing loss of consciousness. All participants were fluent in English, provided informed consent, and received course credit for their involvement. The study was approved by the Florida International University Institutional Review Board.

This study employed a between-subjects design, with 140 participants randomly assigned to one of two experimental groups: a “Face Image Group” presented with images of human faces and an “Object Image Group” presented with images of inanimate objects. Twenty-five participants were excluded from further analysis due to: experimental error (n = 3), inattentiveness during the incidental memory task (n = 2), low accuracy on the Flanker task (below 60%; n = 1), or having fewer than 8 error trials in the Flanker task (n = 19). The final sample included 60 participants in the Object Image Group (*M* = 22.52 years, *SD* = 3.37; 45 female, 15 male) and 55 participants in the Face Image Group (*M* = 22.33 years, *SD* = 3.34; 50 female, 5 male). The sample demographic breakdown reflects typical undergraduate psychology participant pools (Gruber et al., 2021), with a sex imbalance resulting from unbiased enrollment procedures.

### Assessment of SA Symptoms

SA symptoms were assessed using the 7-item Social Anxiety scale from the Screen for Adult Anxiety Related Disorders (SCAARED; Angulo et al., 2017). Participants responded to each item on a 3-point Likert scale (0 = not true or hardly ever true, 1 = somewhat true or sometimes true, 2 = very true or often true), with higher scores indicating greater SA symptom severity. The SCAARED Social Anxiety scale demonstrated adequate internal consistency in the current sample (raw Cronbach’s α = .87).

### Procedure

Participants were randomly assigned to one of two groups, which determined whether the background images in the task were faces or objects. After providing informed consent, participants completed a battery of surveys, including the SCAARED questionnaire. Additional surveys administered during the study examined state/trait assessment of thoughts, feelings, and performance, but are beyond the scope of the current report and not discussed further. Next, participants completed a practice block of a modified Flanker task with unique background images on each trial, requiring an accuracy of at least 75% to proceed; otherwise, the practice block was repeated until this criterion was met. Participants then completed the main Flanker task under peer observation to induce a sense of social evaluation (García Alanis et al., 2019; Niu et al., 2023; Somerville et al., 2013). Subsequently, a surprise incidental memory test was administered using a 2AFC recognition paradigm; each trial presented an image from the Flanker task paired with a novel image of the same type.

### Modified Flanker Task

Participants engaged in a modified Flanker task (Figure 1), where each trial presented an array of five arrows, accompanied by a trial-unique image in the background. Depending on which Image Group participants were assigned to, the background image was either a face or an inanimate object. To prevent potential confounding effects arising from the use of emotionally evocative stimuli, this study utilized neutral face images and neutral inanimate object images, consistent with our previous research employing a similar modified Flanker task (Hosseini et al., 2024). For the Face Image Group, we used the same neutral face dataset created for our previous study (Hosseini et al., 2024), which was developed by merging all existing image sets from the Chicago Face Database (Ma et al., 2015, 2021). Details about the specifics of this dataset can be found in our prior paper (Hosseini et al., 2024). For the Object Image Group, we utilized an internally developed inanimate neutral object dataset. The object images were adjusted to match the face dataset, ensuring that objects were of comparable size, valence, and were centered against a white background with a 2448×1718 resolution (Figure 2). Details regarding dataset development are available in the supplement; the full dataset is accessible via our GitHub repository (https://github.com/NDCLab/mfe-c-dataset). From each dataset, we randomly selected 788 images to be used in the current study.

**Figure 1.** Modified Flanker Task. The figure outlines the sequence of events in a single trial of each version of the Modified Flanker task, as well as the Incidental Memory Task. Image proportions in the figure are adjusted for clarity and do not reflect the actual task display. **(A, C) Modified Flanker Task:** In the Face Image Group (A) and Object Image Group (C), each trial presents a unique background image and fixation rectangle. The trial proceeds as follows: flanking arrows appear, the target arrow appears 150 msecs later, all arrows disappear 200 msecs after the target and the background image remains for the trial duration (jittered 3500-4000 msecs). **(B, D) Incidental Memory Task:** Participants are presented with a fixation cross (jittered 800-1200ms) followed by a self-paced 2AFC recognition test. Note, Figure 1 is available from the authors upon request.

**Figure 2.**
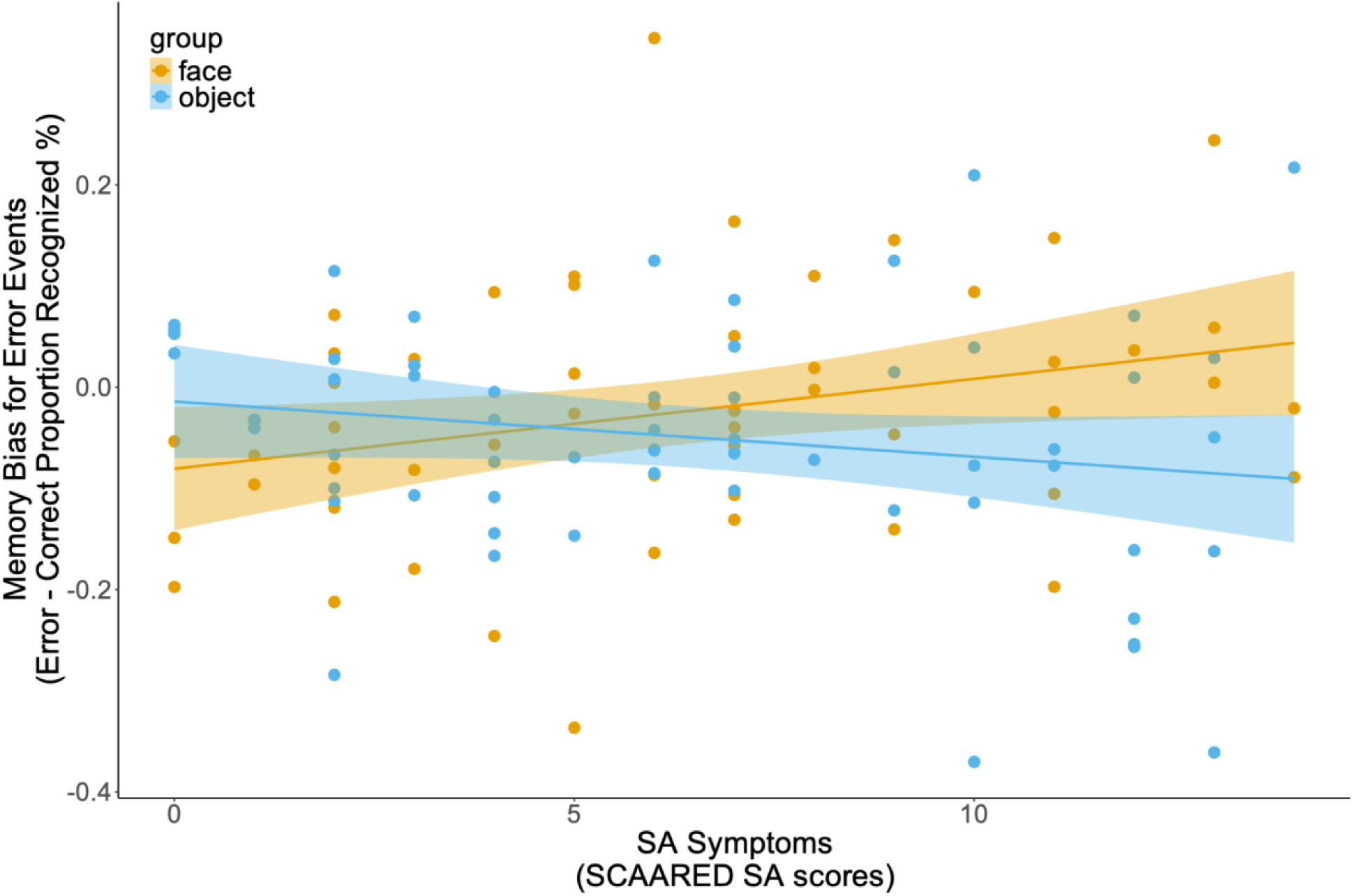
Relationship between SA symptom levels (SCAARED-social scores) and Memory Bias for Error Events in the face and object groups. Higher SA symptom levels, as measured by SCAARED-Social scores, predicted an increased Memory Bias for Error Events in the Face Image Group (*p* = 0.031), but not in the Object Image Group (*p* = 0.14). Raw data are depicted in this figure for ease of interpretation.

Participants indicated the direction (right/left) of a target arrow by key press using their corresponding right/left index fingers; the response window lasted the entire duration of each trial. The target arrow was either congruent (same direction) or incongruent (opposite direction) with the flanking arrows. The arrows appeared for the first 350 msecs of each trial, with the flanking arrows appearing first, followed by the target arrow after a 150 msecs delay. All arrows then remained on screen for 200 msecs before disappearing simultaneously (Figure 1). The background image, with on-screen dimensions of 14.8 cm×10.4 cm, was displayed throughout the duration of each trial. The trial duration randomly jittered between 3500-4000 msecs. The arrow array was surrounded by a fixation rectangle that was centered on the screen and displayed throughout each block. Participants completed 384 trials administered in 12 blocks of equal size. Within each block, congruent and incongruent trials were equally represented. Feedback was provided after each block to maintain sufficient error rates (Gehring et al., 2012). The feedback presented was as follows: “*Good Job*” for accuracy between 75% and 90%, “*Respond Faster*” for accuracy above 90%, and “*Respond More Accurately*” for accuracy below 75%.

Stimuli were presented on a 23.8-inch Dell OptiPlex 7440 AIO computer running Windows 10 with PsychoPy version 2022.2.4 (Peirce et al., 2019). Participants were seated 60 cm from the monitor and used a wired Dell Multimedia Keyboard (KB216) with a US International (QWERTY) layout to record their responses. To ensure participant attentiveness, Flanker trials with reaction times (RTs) faster than 150 msecs (M = 0.04, SD = 0.29) were excluded from subsequent analyses.

### Incidental Memory Task

After the Flanker task, participants completed a self-paced, incidental memory task using a 2AFC approach. Each trial presented participants with two images, one image that previously appeared during the Flanker task, and one new image of the same type (face or object, depending on the assigned participant Image Group). Participants were instructed to select which of the two images they recognized from the previous Flanker task. Participants indicated their choice by pressing a key with their right/left index fingers, corresponding to the position of the image on the screen. Each image pair was displayed until a response was recorded, with each image having on-screen dimensions of 14.8×10.4 cm. Trials were presented in a randomized sequence unique to each participant, and the memory task comprised eight blocks of 48 trials each. The same equipment from the Flanker task was used.

To ensure attentiveness during the memory task, we implemented two exclusion criteria. First, trials (M = 5.53, SD = 28.63) with RTs under 200 msecs were considered too fast and therefore excluded from analyses; participants (n = 2; the same individuals described in the Participants section) were excluded entirely if they had 144 or more such trials (equivalent to three blocks). Second, participants could be excluded for exhibiting repetitive response patterns indicative of inattention across at least three memory blocks (selecting the same answer for at least 20 consecutive trials or for at least 90% of trials within a block); however, no participants were removed due to this second criterion.

### Memory Bias for Error Events

Consistent with our previous work (Hosseini et al., 2024), we computed a “Memory Bias for Error Events” score, capturing the difference in recognition accuracy for images that originally appeared during error vs. correct responses in the Flanker task. Specifically, we first calculated the percentage of correctly recognized images (“proportion recognized”), separately for images that originally appeared on error and correct trials during the Flanker task (equations 1 and 2). We then computed a difference score by subtracting the proportion recognized for images that originally appeared on correct trials from the proportion recognized for images that originally appeared on error trials (Equation 3). The resulting difference score indicates the relative incidental memory performance for error-related images, which we refer to as Memory Bias for Error Events, and which was used in subsequent analyses. To control for stimulus congruency and isolate on error-related effects, we calculated the proportion recognized using only images from incongruent-error and incongruent-correct trials (Buzzell et al., 2019; Eriksen & Eriksen, 1974). Hereafter, for simplicity, the terms “error” and “correct” trials are used in place of incongruent-error and incongruent-correct trials, respectively.

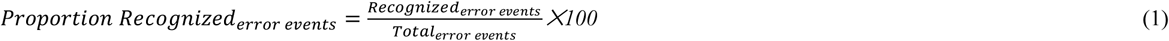

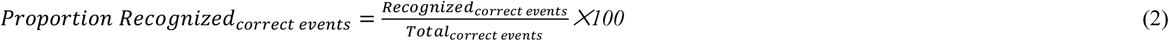

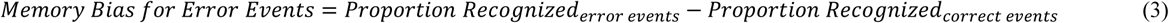

### Analytic Plan

We investigated the influence of Flanker task performance on subsequent recognition memory, and how this relationship might differ based on the nature of stimuli and individual differences in SA symptom levels. We first conducted preliminary checks to confirm the presence of typical congruency effects (“Flanker effects”) in our modified Flanker tasks, assessing both accuracy and reaction times. Next, we performed preliminary analyses to explore whether Flanker task accuracy (correct vs. error trials) was predictive of subsequent recognition memory performance, irrespective of individual differences in SA symptom levels. Finally, our primary analyses (hypothesis testing) focused on whether SA symptom levels were predictive of Memory Bias for Error Events, and whether this effect was specific to social stimuli (faces) or extended to nonsocial stimuli (objects). Statistical analyses were conducted in R (version 4.4.1; R Core Team, 2022) using the lme4 (version 1.1-36; Bates et al., 2015), glmmTMB (version 1.1.10; Brooks et al., 2017), and lmerTest (version 3.1-3; Kuznetsova et al., 2017) packages, as well as the lm function from the stats package (version 4.4.1); further details can be found in the supplement.

**Preliminary Analyses.** To confirm the presence of typical Flanker congruency effects (Eriksen & Eriksen, 1974), we analyzed Flanker accuracy and response time (RT) as a function of congruency. We employed a mixed-effects model approach, given that our outcomes were repeated measures and to address issues of non-normality and heteroscedasticity. Specifically, a generalized linear mixed model (GLMM; equation 4) was employed for accuracy data, and a linear mixed-effects model (LMM; equation 5) was adopted for analyses of RT. Both models tested for congruency effects overall, as well as potential task-related differences between the Face and Object Image groups. Additionally, to rule out the possibility that Flanker task performance differed as a function of SA symptom levels, we re-ran these models when adding interaction terms for SA symptom levels (see supplement). Finally, a preliminary LMM (equation 6) was also fit to explore whether Flanker task accuracy (error vs. correct) was predictive of subsequent recognition memory performance overall, regardless of SA symptom levels, and whether any potential effect differed as a function of the Flanker task version (Face vs. Object Image Group). In all models, a random intercept was included for participants. Further statistical details for these models can be found in the Supplement.

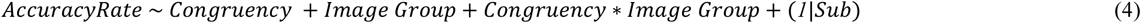

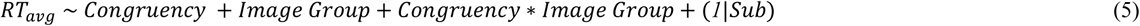

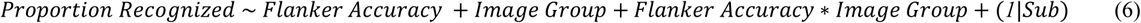

**Hypothesis Testing.** Our primary analyses served to achieve two objectives. First, to test the hypothesis that SA symptom levels predict Memory Bias for Error Events (the proportion recognized difference score; equation 3), replicating the results of our prior study (Hosseini et al., 2024) that employed face (social) stimuli. Second, to test whether the hypothesized relationship between SA symptom levels and Memory Bias for Error Events is specific to social stimuli (faces) as opposed to reflecting a more general tendency to encode any information (objects) present during error events. To this end, we simultaneously carried out both of these tests by fitting a single linear regression model (equation 7) predicting Memory Bias for Error Events (equation 3) from SA symptom, Image Group (face vs. object), and their interaction. Simple linear regression was employed for this analysis, given that a mixed modeling approach would be redundant in the absence of a repeated measures outcome.

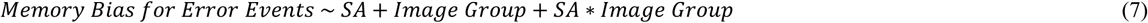

## Results

### Preliminary Results

Confirming the validity of our Flanker task paradigms, mixed-effects analyses resulted in the expected congruency effects for both accuracy and RT, consistent with a classic Flanker effect (Eriksen & Eriksen, 1974). Specifically, the GLMM (equation 4) predicting accuracy (see Supplemental Table S1) yielded a significant main effect of congruency (standardized *β* = –1.987, standardized 95% CI [-2.257, –1.716], *z* = –14.416, *p* < .001), with accuracy being lower on incongruent vs. congruent trials. Similarly, the LMM (equation 5) predicting RT (see Supplemental Table S2) identified a significant main effect of congruency (standardized *β =* 0.763, standardized 95% CI [0.673, 0.852], *t*(111.7) = 16.872, *p* < .001), with RTs being slower on incongruent vs. congruent trials. No further significant main effects of Image Group, nor interactions between Image Group and congruency were observed for either accuracy or RT (all *p* > .165). Thus, analyses of Flanker task behavior confirmed the presence of expected congruency effects, and further demonstrated that participants exhibited similar overall performance for both versions of the flanker task (face, object).

Before testing our primary hypotheses regarding associations between SA symptoms and Memory Bias for Error Events, we explored whether recognition memory performance overall differed as a function of Flanker task accuracy (error vs. correct), task (Face vs. Object Image Group), or their interaction. The LMM testing these relations (equation 6; Supplemental Table S3) revealed a significant main effect of Image Group, such that objects were remembered better than faces (standardized *β =* 0.809, standardized 95% CI [0.467, 1.151], *t*(205.14) = 4.663, *p* < .001). No further main effects or interactions were identified in this model (all *p* > .104).

### SA-Related Memory Bias for Error Events is Specific to Faces

Primary hypothesis testing was carried out via linear regression analysis to examine whether SA symptom levels predicted Memory Biases for Error Events, and whether this relationship was specific to social stimuli (Face Image Group) or reflected a more general tendency to encode any information present during errors events (Object Image Group). This regression analysis revealed a significant interaction between SA symptom levels and group (Face vs. Object Image Group) in predicting Memory Bias for Error Events (standardized *β =* –0.483, standardized 95% CI [-0.85, –0.117], *t*(110) = –2.613, *p* = 0.01); see Figure 2 and Table 1. To understand the nature of this interaction, simple slopes analyses were performed. In the Face Image Group, higher SA symptom levels significantly predicted an increased Memory Bias for Error Events, as evidenced by better recognition of faces that previously appeared during error vs. correct trials (standardized *β* = 0.298, standardized 95% CI [0.027, 0.569], *t*(110) = 2.179, *p* = 0.031). In contrast, the Object Image Group exhibited no such association between SA symptom levels and Memory Bias for Error Events (standardized *β* = –0.183, standardized 95% CI [-0.428, 0.061], *t*(110) = –1.485, *p* = 0.14). Therefore, the data indicate that the enhanced Memory Bias for Error Events in individuals with high SA symptoms is specific to social stimuli (faces) and does not extend to non-social stimuli (objects) for the current paradigm. See Table 1 for full results.

**Table 1.**
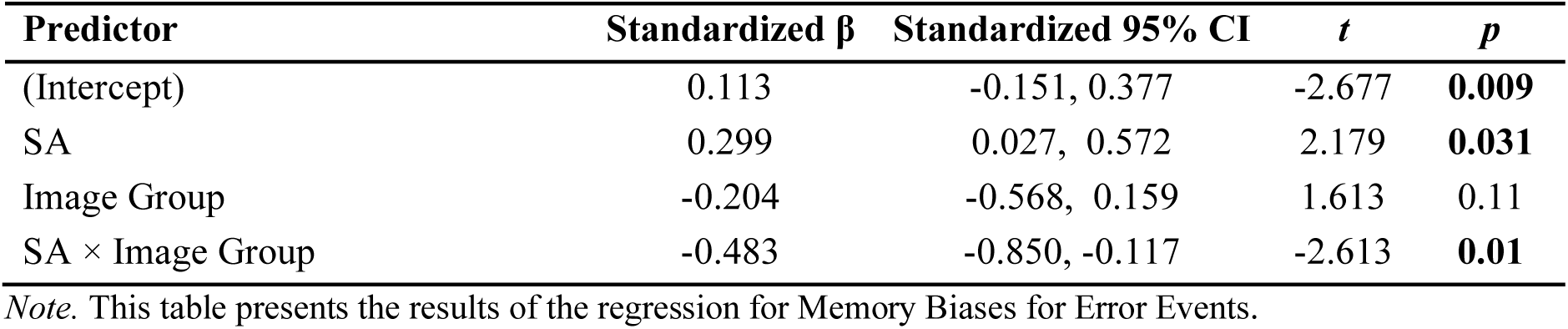
Summary of the regression Analysis Predicting Memory Biases for Error Events.

| Predictor | Standardized $\beta$ | Standardized 95% CI | $t$ | $p$ |
| --- | --- | --- | --- | --- |
| (Intercept) | 0.113 | -0.151, 0.377 | -2.677 | **0.009** |
| SA | 0.299 | 0.027, 0.572 | 2.179 | **0.031** |
| Image Group | -0.204 | -0.568, 0.159 | 1.613 | 0.11 |
| SA × Image Group | -0.483 | -0.850, -0.117 | -2.613 | **0.01** |

*Note.* This table presents the results of the regression for Memory Biases for Error Events.

### Ruling Out Alternative Interpretations

Our memory bias results are consistent with the interpretation that, for individuals high in SA symptoms, the commission of errors leads to enhanced encoding of social-specific information present during error events. However, an alternative interpretation is that individuals high in SA symptoms are more distracted by (and pay more attention to) social background images to begin with, which then causes errors to occur on the Flanker task. However, if this alternative distraction-based account were correct, then individuals high in SA should exhibit worse Flanker task performance for the social Flanker task (i.e., slower and less accurate responses due to distraction by the background images). To test this alternative hypothesis, we re-ran our mixed-effects analyses predicting Flanker task accuracy and RT (see “Preliminary Results”) when further including interaction terms for SA symptoms. Crucially, results of these models yielded no significant main effects or interactions involving SA symptoms in the prediction of RT or accuracy (all *p* > .14); see Supplemental Tables S4 and S5). These results provide no support for a distraction-based alternative explanation of the association between SA symptom levels and Memory Bias for Error Events.

## Discussion

The present study had two primary aims. First, within a larger sample and using an improved memory assessment approach, we sought to replicate our prior finding that individuals with higher SA symptoms exhibit enhanced memory for information present during error events (i.e., Memory Bias for Error Events, (Hosseini et al., 2024). Second, we tested the hypothesis that this SA-related memory bias would be specific to socially-relevant stimuli (i.e., faces) rather than reflecting a general tendency to encode any information present during error events (e.g, objects). Replicating our prior findings, for individuals that completed the modified Flanker task involving background face stimuli, higher SA symptoms predicted an enhanced Memory Bias for Error Events. Moreover, as hypothesized, a significant association between Memory Bias for Error Events and SA symptom levels was only present for individuals that completed the Flanker task involving social stimuli (faces). In other words, higher SA symptom levels predicted better memory for face images that had originally appeared during error (vs. correct) trials, but no such relationship emerged for object images. These findings provide evidence that error-related memory biases in SA do not reflect a non-specific enhancement of information encoding when errors occur, but instead, reflect the selective encoding of socially-relevant stimuli present during error events.

The findings provide a robust replication of the association between SA symptoms and Memory Bias for Error Events, which was originally reported in our prior study (Hosseini et al., 2024). In the current investigation, we improved upon the limitations of our prior study by employing a larger participant sample and, importantly, by replacing the old/new recognition paradigm used in our prior work with a 2AFC task. A 2AFC design requires making a relative judgment between two stimuli on each trial and further balances the ratio of presented old/new stimuli (each trial presents one old and one new image). As a result, a 2AFC design is less susceptible to a potential confound of response bias (i.e., individual differences in the overall tendency to respond “old” vs. “new”; (Kellen et al., 2018; Kroll et al., 2002; Van Zandt, 2000). Thus, the methodological improvements of the current study, coupled with the consistency of the observed effect across two different recognition paradigms in two different studies, increases confidence that the observed association between SA symptoms and Memory Bias for Error Events reflects a true effect. Accordingly, these data further establish a robust methodological foundation that can be leveraged by future studies to examine how error-related social cognition, through memory, may ultimately contribute to SA maintenance.

Beyond replication, the central contribution of this study is the demonstration that relations between SA symptoms and an enhanced Memory Bias for Error Events is specific to social stimuli. Consistent with our hypothesis, we suggest that the social specificity of this effect is a result of the greater motivational relevance of social stimuli for individuals high in SA, combined with the notion that errors are salient events that trigger increased arousal and attention towards motivationally relevant stimuli. Prior work demonstrates that other kinds of salient events lead to biased processing of motivationally relevant stimuli present at the time of the event, resulting in biased encoding (Clewett et al., 2018; Dunsmoor et al., 2015; Hutmacher & Kuhbandner, 2020; Lin et al., 2010; Patil et al., 2017; Swallow et al., 2022; Swallow & Jiang, 2010, 2011). Biased encoding in response to salient events is thought to arise as the result of NE release driving increased neural activity within brain areas concurrently active due to their processing of information that is either task relevant or otherwise motivationally relevant to the individual, and suppression of other neural activity (i.e., increased “neuronal gain”; Dahl et al., 2022; De Martino et al., 2008; Mather et al., 2016; Mather & Sutherland, 2011; Ullsperger et al., 2010; Wessel, 2018). Although evidence that errors elicit NE release is indirect (for a review, see: Wessel, 2018), direct evidence for NE release following other kinds of salient events have been reported (Mather et al., 2016; Stanley et al., 2023; Vazey et al., 2018; Ventura et al., 2008; Wilson et al., 2023). Moreover, it has been documented that both errors and other salient events elicit an increase in attention and arousal (Compton et al., 2021; Steinhauser & Andersen, 2019; Weinbach & Henik, 2014). Importantly, for individuals with SA, whose core fear is negative social evaluation (Clark & Wells, 1995; Rapee & Heimberg, 1997; Zhang et al., 2023), social cues like faces are motivationally relevant (Cooney et al., 2006; Kivity & Huppert, 2016), particularly within the context of having made a performance failure. Therefore, when an individual high in SA commits an error, we suggest that this results in biased processing of social information, leading to the selective memory enhancement for concurrently presented faces—but not objects—observed in our data. Broadly, this interpretation aligns with findings that cognitive biases in SA are tuned specifically to social-evaluative information (e.g., Romano et al., 2020). However, future work that incorporates proximal measures of NE release following error commission (e.g., pupil dilation) could provide further tests of the underlying mechanism that leads to the observed behavioral phenomenon reported here.

The present findings have important implications for both theory and treatment of SA. Our results provide the first empirical evidence that the Memory Bias for Error Events in SA is socially specific, supporting cognitive models that emphasize the role of biased processing of social-evaluative information in maintaining SA symptoms (Clark & Wells, 1995; Rapee & Heimberg, 1997). This social specificity suggests that when individuals high in SA symptoms make errors, they do not simply exhibit enhanced encoding of all error-related information, instead, they selectively attend to and remember social cues during moments of perceived failure. Although we were unable to assess causality in the current study, these findings are consistent with the notion that enhanced error monitoring in SA (Barker et al., 2015; Endrass et al., 2014; Hosseini et al., 2024; Judah et al., 2016; Kujawa et al., 2016; Meyer et al., 2021; Niu et al., 2023) may ultimately play a causal role in the maintenance and worsening of SA. That is, in line with cognitive models of SA (Clark & Wells, 1995; Rapee & Heimberg, 1997), it is possible that enhanced error monitoring in SA drives a stronger Memory Bias for Error Events, which in turn strengthens negative beliefs about one’s abilities within social domains and ultimately maintains or worsens SA (Hosseini et al., 2024). However, further research employing longitudinal assessment is needed to directly test such a causal hypothesis. From a clinical perspective, these findings have the potential to ultimately inform the development of more targeted intervention strategies for individuals with SA. With further validation in clinical samples and via longitudinal assessment, treatments might benefit from targeting the selective encoding of social information during error events, rather than addressing error monitoring or memory biases more broadly. Such approaches could potentially involve techniques that reduce the salience of social cues during performance situations or that modify the interpretation of social information when errors occur—though additional research would be needed to develop and test such interventions.

Additional analyses ruled out alternative explanations of the observed relations between SA symptoms and Memory Bias for Error Events. For example, one alternative interpretation is that the association between SA symptoms and memory for faces from error trials arises not due to error commission driving attention toward face stimuli, but instead, that faces are more distracting for individuals high in SA to begin with, drawing attention away from the flanker task and leading to errors. However, our data do not support such a distraction-based alternative interpretation. If distraction by face stimuli were the primary mechanism, then higher SA symptoms should have been associated with worse Flanker task performance (i.e., lower accuracy or slower RTs), particularly for individuals that performed the task with face images. In contrast, our analyses revealed no relationship between SA symptom levels and either Flanker task accuracy or RT. These behavioral analyses, combined with our previous work showing no link between stimulus-evoked neural responses to faces and SA symptoms or memory bias levels (Hosseini et al., 2024), argue strongly against a distraction-based account. Instead, the evidence supports the interpretation that an enhanced Memory Bias for Error Events in high SA individuals arises as the *result* of error commission. Further evidence supporting this interpretation also comes from our prior study, which demonstrated that only response-related neural activity on error trials (i.e. activity associated with error monitoring) predicted the subsequent effect of Memory Bias for Error Events (Hosseini et al., 2024).

Orthogonal to our primary effects of interest, an unexpected finding emerged from the analysis of overall recognition memory performance, irrespective of trial accuracy (correct vs. error). We observed a significant main effect of Image Group, indicating that participants demonstrated better incidental recognition memory for objects compared to faces (approximately 9% higher proportion recognized for objects). This observation contrasts with some previous research, often employing intentional learning paradigms, which reported superior memory for faces over other stimulus categories (Dobson & Rust, 1994; Sato & Yoshikawa, 2013). However, the divergence in our findings might stem from key methodological differences, including our focus on incidental rather than intentional memory encoding, or potentially our use of a larger stimulus set. Importantly, the main effect of Image Group on overall memory performance does not alter the interpretation of our primary finding: the significant interaction effect showing that the relationship between SA symptoms and Memory Bias for Error Events was specifically present for social (face) stimuli and absent for non-social (object) stimuli.

### Limitations and future directions

The present study provides compelling evidence for the social specificity of an enhanced Memory Bias for Error Events in individuals with high SA symptoms. However, as no study is without limitations, we highlight important avenues for future research. The current study assessed associations with SA symptoms within an unselected undergraduate sample. Therefore, studies testing these associations among clinically-referred participants with social anxiety disorder are needed to assess generalizability of the findings to clinical populations. Relatedly, the unbiased enrollment procedures employed in the current study resulted in a sex imbalance among participants (consistent with the demographics of the undergraduate psychology student population). Future research could employ targeted recruitment strategies to achieve a sex-balanced sample and assess generalizability of findings. Given the cross-sectional nature of the current study, causality cannot be assessed and future longitudinal assessments are needed. The current study establishes an important boundary condition for the association between SA symptoms and Biased Memory for Error Events, demonstrating that this effect is specific to social stimuli. However, it remains to be tested whether this effect is also specific to instances of self-directed errors, as studied here, or if it would generalize to instances of externally-presented error/negative feedback as well. We hypothesize that the Memory Bias for Error Events effect would generalize to externally–presented feedback, and given the high relevance of negative external feedback to the interpersonal fears inherent in SA, future work that directly tests this hypothesis would be particularly informative. Similarly, studies that directly manipulate the emotional valence of background social stimuli would be informative. Given that a core fear in SA is negative social evaluation (Clark & Wells, 1995; Rapee & Heimberg, 1997; Zhang et al., 2023), it is plausible that the Memory Bias for Error Events in high SA individuals would be even more pronounced for faces displaying negative emotions (e.g., anger, disgust). Conversely, it is an open question as to whether positive social cues (e.g., happy faces) would exhibit similar effects, or possibly even buffer against such biases. Additional clarification of these boundary conditions could further refine theory and inform the development of targeted intervention approaches.

## Conclusion

This study replicates prior work by demonstrating that individuals with higher SA symptoms exhibit an enhanced Memory Bias for Error Events, and crucially, establishes that this bias is specific to social stimuli (faces) rather than reflecting a general tendency to encode any information present during errors. The absence of this memory bias for non-social stimuli (objects) reveals a critical boundary condition that underscores the importance of social-evaluative concerns in SA-related cognitive processes. These results advance theoretical understanding by providing the first empirical evidence for social specificity in error-related memory biases, supporting cognitive models that emphasize biased processing of socially-relevant information in SA. If further substantiated in clinical populations and via longitudinal methods, these findings suggest that interventions for SA may be most effective when they specifically target the selective encoding of social information during error events rather than addressing error monitoring or memory biases broadly.

## Declarations

### Author contributions

**Conceptualization**: K. Hosseini, J.W. Pettit, A.T. Mattfeld, and G.A. Buzzell; **Methodology**: K. Hosseini and G.A. Buzzell; **Formal Analysis**: K. Hosseini; **Investigation**: K. Hosseini; **Data Curation**: K. Hosseini; Software: K. Hosseini; **Validation**: K. Hosseini and G.A. Buzzell; **Visualization**: K. Hosseini; **Writing – Original Draft Preparation**: K. Hosseini; **Writing – Review & Editing**: K. Hosseini, J.W. Pettit, A.T. Mattfeld, and G.A. Buzzell; **Supervision**: G.A. Buzzell; **Project Administration**: K. Hosseini and G.A. Buzzell; **Resources**: G.A. Buzzell; **Funding Acquisition**: J.W. Pettit and G.A. Buzzell.

## Acknowledgments

We would like to thank all undergraduate research assistants at the Neural Dynamics of Control Lab that assisted with data collection. We also thank the participants taking part in the study.

## Supplemental Material

The supplemental materials detail the object image dataset development, statistical analysis overview, preliminary results tables (Tables S1-S3), and additional analyses to rule out SA-related variability in Flanker task performance (Tables S4-S5).

## Prior versions

A preprint of this manuscript was published on bioRxiv (add the link here).

