## Supplemental Materials for "Error-Related Memory Biases Are Specific to Social Stimuli for Socially Anxious Individuals"

### Object Image Dataset Development

The object image dataset was developed to be used in the Object Image Group, as reported in the main text. Here, the primary goal was to create a set of neutral, inanimate object images that were comparable in visual properties to the face stimuli used in the Face Image Group.

The development process consisted of several stages:

1. Image Sourcing: A large pool of high-resolution images was compiled using the DuckDuckGo, Google, and Bing search engines, as well as using stock photography databases such as Pexels.com. Search queries were restricted to neutral, everyday items (e.g., "chair," "lamp," "car," "book," "cup"). All selected images were verified to be licensed as "free to use" to avoid copyright restrictions, and the highest possible quality version of each image was downloaded.
2. Selection and Curation: From the initial pool of images obtained, two independent raters curated the images based on a comprehensive set of predefined criteria to ensure neutrality, consistency, and high image quality:
    - Neutral Valence: Objects related to war, military weapons, or any aggressive-looking tools were excluded. Similarly, objects were required to be in a normal, ordinary condition (e.g., an image of a wrecked car was deemed not acceptable).

- Selection Scope: Images were selected from a diverse collection of objects, spanning multiple categories from inanimate, natural items to constructed objects. Object images deemed "abstract", "exotic" or otherwise "out of the ordinary" were excluded.
- Visual Distinctiveness: All images in the final image set were required to be deemed visually distinct from one another. While multiple images of the same object category were permissible, they had to be clearly distinguishable (e.g., reflecting a different viewing angle and a different color/design). Images of the same object from only a different angle were deemed not sufficient.
- Composition and Focus: Each image was required to feature a single, centrally-located object. The full object was preferred, without being cropped or partially shown. Images containing multiple prominent objects were avoided.
- Image Quality: All images were required to be high-resolution, colored (not grayscale), and free of visual defects. Images that were blurry, pixelated, taken in low-light conditions, or captured from an unusual perspective were excluded. The objects depicted had to be real photographs, not drawings.
- Exclusion of Confounding Elements: Images were excluded if they contained human or animal figures (or parts thereof), offensive or inappropriate content, watermarks, or prominent text or logos. Small, incidental brand names that were not easily legible were considered acceptable.
- Background: A homogeneous background (e.g., uniform in color and texture) was strongly preferred to simplify background removal (final images were converted to have a solid white background).

3. Image Processing: All selected images were processed to match the specifications of the Chicago Face Database stimuli used in the Face Image Group. Thus, the background of each image was removed and substituted with a uniform white (#FFFFFF) background. This was accomplished using the Google-developed background removal tool accessible via Google Drawings. Then, images were resized and cropped to a standard dimension of 2448 × 1718 pixels to match the face dataset. The object was centered within the frame.

A final set of 788 unique object images was used in the modified Flanker task and the subsequent incidental memory assessment. This object image dataset is accessible via our GitHub repository, and we encourage reuse by other researchers: https://github.com/NDCLab/mfe-c-dataset. For reuse, please cite the current manuscript, as well as the GITHUB repository.

**Statistical Analysis Overview**

Statistical analyses used R (version 4.4.1; R Core Team, 2022). Two Linear Mixed-effects Models (LMMs) were run using the lme4 package (version 1.1-36; Bates et al., 2015), and a Generalized Linear Mixed Model (GLMM) was run using the glmmTMB package (version 1.1.10; Brooks et al., 2017). All the LMMs and GLMM included random intercepts for participants. The lmerTest package (version 3.1-3; Kuznetsova et al., 2017) was used for LMM significance calculations, employing Satterthwaite's method for estimating degrees of freedom and generating p-values. Further, a linear regression was performed using the lm function in the stats package (version 4.4.1). The LMM and linear regression assumptions were examined. If any of the assumptions were violated, an alternative approach (e.g., GLMM) was adopted and noted in subsequent sections. Standardized coefficients and confidence intervals are reported throughout.

**Analysis of Flanker Task Performance**

To assess the typical congruency effects in our modified Flanker tasks (Eriksen & Eriksen, 1974), we conducted two analyses. First, we examined how Flanker congruency influenced accuracy using a Generalized Linear Mixed Model (GLMM; equation 1). This model was selected after residual plots revealed a non-normal distribution and heteroscedasticity in the accuracy data. The GLMM with a beta distribution and a logit link function was employed as it is particularly suitable for analyzing proportional data such as accuracy rates (Chen et al., 2017; Cribari-Neto & Zeileis, 2010). Second, we investigated the impact of Flanker congruency on reaction times (RTs) using a Linear Mixed-effects Model (LMM; equation 2). A random intercept for participants was included in both models. Both models examined the basic congruency effects while also testing for potential variations across Image Groups (Face vs. Object). Only correct trials were used to compute the mean RTs for RT analysis.

$$AccuracyRate \sim Congruency \ + Image\ Group + Congruency * Image\ Group + (1|Sub) \qquad (1)$$

$$RT_{avg} \sim Congruency \ + Image\ Group + Congruency * Image\ Group + (1|Sub) \qquad (2)$$

**Preliminary Results Tables**

**Table S1**

*Summary of the GLMM Analysis Predicting Flanker Task Accuracy*

| **Predictor** | **Standardized β** | **Standardized 95% CI** | ***z*** | ***p*** |
|---|---|---|---|---|
| (Intercept) | 3.694 | 3.417, 3.972 | 26.126 | ***p* < .001** |
| Flanker Congruency | -1.987 | -2.257, -1.716 | -14.416 | ***p* < .001** |
| Image Group | -0.189 | -0.549, 0.170 | -1.033 | 0.302 |
| Flanker Congruency × Image Group | 0.098 | -0.262, 0.457 | 0.532 | 0.594 |

*Note.* This table presents the results of the GLMM for accuracy in the Flanker task.

**Table S2**

*Summary of the LMM Analysis Predicting Flanker Task RT*

| Predictor | Standardized β | Standardized 95% CI | *t* | *p* |
|---|---|---|---|---|
| (Intercept) | -0.344 | -0.598, -0.089 | 44.835 | ***p* < .001** |
| Flanker Congruency | 0.763 | 0.673, 0.852 | 16.872 | ***p* < .001** |
| Image Group | -0.004 | -0.356, 0.348 | -0.024 | 0.981 |
| Flanker Congruency × Image Group | -0.087 | -0.209, 0.036 | -1.390 | 0.166 |

*Note.* This table presents the results of the LMM for RT in the Flanker task.

**Table S3**

*Summary of the LMM Analysis Predicting Proportion Recognized*

| Predictor | Standardized β | Standardized 95% CI | *t* | *p* |
|---|---|---|---|---|
| (Intercept) | -0.255 | -0.503, -0.007 | 40.829 | ***p* < .001** |
| Flanker Accuracy | -0.243 | -0.537, 0.051 | -1.626 | 0.105 |
| Image Group | 0.809 | 0.467, 1.151 | 4.663 | ***p* < .001** |
| Flanker Congruency × Image Group | -0.201 | -0.606, 0.204 | -0.977 | 0.330 |

*Note.* This table presents the results of the LMM for Proportion Recognized in the Flanker task.

### Ruling Out SA-Related Variability in Flanker Task Performance

To rule out the possibility that Flanker task performance differed as a function of SA symptom levels, we re-ran the mixed-effects models testing the Flanker effects (model 1 and 2) when adding interaction terms for SA symptom levels (equations 3 and 4).

$$AccuracyRate \sim Congruency \ + Image\ Group + SA + Congruency * Image\ Group * SA + (1|Sub) \quad (3)$$

$$RT_{avg} \sim Congruency \ + Image\ Group + SA + Congruency * Image\ Group * SA + (1|Sub) \quad (4)$$

Results from these expanded models revealed no significant main effects or interactions involving SA symptoms in the prediction of either accuracy (Table S4) or RT (Table S5). These findings provide no support for the alternative hypothesis that SA symptom levels influence Flanker task performance, thereby ruling out a distraction-based explanation for the observed Memory Bias for Error Events.

**Table S4**

*Summary of the GLMM Analysis Predicting Flanker Task Accuracy with the inclusion of SA in the Model*

| Predictor | Standardized β | Standardized 95% CI | *z* | *p* |
|---|---|---|---|---|
| (Intercept) | 3.878 | 3.354, 4.403 | 14.492 | ***p* < .001** |
| Flanker Congruency | -2.183 | -2.702, -1.663 | -8.234 | ***p* < .001** |
| Image Group | -0.363 | -1.045, -0.320 | -1.041 | 0.298 |
| SA | -0.029 | -0.098, 0.039 | -0.832 | 0.406 |
| Flanker Congruency × Image Group | 0.378 | -0.302, 1.058 | 1.089 | 0.276 |
| Flanker Congruency × SA | 0.031 | -0.037, 0.099 | 0.89 | 0.374 |
| Image Group × SA | 0.028 | -0.064, 0.120 | 0.599 | 0.549 |
| Flanker Congruency × Image Group × SA | -0.045 | -0.136, 0.047 | -0.958 | 0.338 |

*Note.* This table presents the results of the GLMM for accuracy in the flanker task with SA symptom levels included in the model.

**Table S5**

*Summary of the LMM Analysis Predicting Flanker Task RT with the inclusion of SA in the Model*

| Predictor | Standardized β | Standardized 95% CI | *t* | *p* |
|---|---|---|---|---|
| (Intercept) | -0.349 | -0.603, -0.095 | 24.932 | ***p* < .001** |
| Flanker Congruency | 0.763 | 0.673, 0.853 | 8.189 | ***p* < .001** |
| Image Group | 0.002 | -0.349, 0.354 | -0.441 | 0.659 |
| SA | -0.192 | -0.454, 0.070 | -1.445 | 0.150 |
| Flanker Congruency × Image Group | -0.087 | -0.211, -0.037 | -0.634 | 0.527 |
| Flanker Congruency × SA | 0.025 | -0.069, 0.119 | 0.516 | 0.606 |
| Image Group × SA | 0.094 | -0.258, 0.447 | 0.526 | 0.599 |
| Flanker Congruency × Image Group × SA | -0.007 | -0.132, 0.118 | -0.114 | 0.910 |

*Note.* This table presents the results of the LMM for RT in the Flanker task with SA symptom levels included in the model.

### Participant Demographics

**Table S6**

*Participant Demographics: Sex at Birth, Race/Ethnicity, and Education*

| Sex (at Birth) | Count | % |
|---|---|---|
| Female | 113 | 80.71 |
| Male | 27 | 19.28 |

| **Race/Ethnicity** | | |
|---|---|---|
| Hispanic/Latino/a/x | 88 | 62.86 |
| Black | 18 | 12.86 |
| White | 5 | 3.57 |
| Asian | 3 | 2.14 |
| Hispanic/Latino/a/x; White | 17 | 12.14 |
| Black; Hispanic/Latino/a/x | 4 | 2.86 |
| Black; White | 1 | 0.71 |
| Hispanic/Latino/a/x; Middle Eastern/N. African | 3 | 2.14 |
| Asian; Black; Hispanic/Latino/a/x | 1 | 0.71 |
| **Education** | | |
| Master's degree | 1 | 0.71 |
| Some post-undergrad work | 1 | 0.71 |
| Bachelor's degree | 18 | 12.86 |
| Associate's degree | 79 | 56.43 |
| Some college | 30 | 21.43 |
| High school diploma | 11 | 7.86 |

*Note.* This table displays the demographic information of participants enrolled in the study prior to any exclusions.
